# EZH2 co-opts gain-of-function p53 mutants to promote cancer growth and metastasis

**DOI:** 10.1101/298737

**Authors:** Yu Zhao, Liya Ding, Dejie Wang, Zhenqing Ye, Yunqian Pan, Linlin Ma, Roger Coleman, Ilsa Coleman, R. Jeffery Karnes, Jun Zhang, Peter S. Nelson, Liguo Wang, Runzhi Zhu, Haojie Huang

## Abstract

With the unfolding of more and more cancer-driven gain-of-function (GOF) mutants of p53, it is important to define a common mechanism to systematically target different mutants rather than develop strategies tailored to inhibit each mutant individually. Here, using RNA immunoprecipitation sequencing (RIP-seq) we identified EZH2 as a p53 mRNA-binding protein. EZH2 bound to the internal ribosome entry site (IRES) in the 5’ untranslated region (5’UTR) of p53 mRNA and enhanced p53 protein translation in a methyltransferase-independent manner. EZH2 augmented p53 GOF mutant-mediated cancer growth and metastasis by increasing p53 GOF mutant protein level. EZH2 overexpression associated with the worse outcome only in patients with p53-mutated cancer. Depletion of EZH2 by antisense oligonucleotides inhibited p53 GOF mutant-mediated cancer growth. Our findings reveal a non-methyltransferase function of EZH2 that controls protein translation of p53 GOF mutants, inhibition of which causes synthetic lethality in cancer cells expressing p53 GOF mutants.

## INTRODUCTION

EZH2 is a SET domain-containing protein that belongs to the Polycomb group (PcG) family (Margueron & Reinberg, 2011). Working coordinately with other PcG proteins in the Polycomb repressive complex 2 (PRC2), EZH2 primarily functions as an methyltransferase by catalyzing histone H3 lysine 27 trimethylation (H3K27me3) (Cao *et al*, 2002, Czermin *et al*, 2002, Kuzmichev *et al*, 2002, Muller *et al*, 2002). The Polycomb-dependent (PcD) function of EZH2 is not only important for developmental patterning, X-chromosome inactivation, stem cell maintenance and cell-fate decision (Boyer *et al*, 2005, Ezhkova *et al*, 2009, Margueron & Reinberg, 2011, Plath *et al*, 2003), but also implicated in cancer (Cha *et al*, 2005, Chen *et al*, 2010, Varambally *et al*, 2002). Overexpression of EZH2 often correlates with aggressive, metastatic solid tumors such as prostate and breast cancers (Karanikolas *et al*, 2009, Kleer *et al*, 2003, Varambally *et al*, 2002). Active mutations in the SET domain of EZH2 frequently occur in human lymphomas (McCabe *et al*, 2012a, Morin *et al*, 2010, Sneeringer *et al*, 2010), resulting in aberrant activation of PcD and elevation of H3K27me3.

In a Polycomb-independent (PcI) manner, EZH2 acts in “solo” to regulate actin polymerization and the oncogenic activities of transcription factors such as the androgen receptor (AR), but the PcI function remains methyltransferase-dependent (Su *et al*, 2005, Xu *et al*, 2012). Because of the importance of the methyltransferase-dependent PcD and PcI functions in cancer, targeting the enzymatic activity of EZH2 is the current focus to develop small molecule inhibitors of EZH2 for cancer treatment (McCabe *et al*, 2012b). A few such inhibitors have been developed and exhibit apparent anti-cancer activities by decreasing cell proliferation and tumor growth in various cancer models (McCabe *et al*, 2012b, Wu *et al*, 2016). While the methyltransferase activities of EZH2 are well studied, it remains unexplored whether or not EZH2 also possesses non-methyltransferase function(s) that might also be important for oncogenesis.

*TP53* is a well-studied tumor suppressor gene (Levine, 1997, Li *et al*, 2012). It is commonly mutated in advanced tumors. While losing the tumor suppressor activity, some mutants of p53 acquire a dominant negative function to inhibit the activity of the remaining wild-type p53 or gain completely new functions (GOF) to drive cancer progression, which include the functions to promote cell proliferation, migration, metastasis, and metabolism in various types of cancer (Dittmer *et al*, 1993, Freed-Pastor *et al*, 2012, Olive *et al*, 2004, Weissmueller *et al*, 2014). Due to the presence of many different types of p53 GOF mutations with distinctive roles in driving cancer progression, various strategies have been explored to target mutant p53 for cancer therapy, including the degradation of p53 mutant proteins, conversion back to the wild-type p53 or targeting downstream signaling pathways of p53 mutants (Adorno *et al*, 2009, Muller & Vousden, 2014, Weissmueller *et al*, 2014, Zhu *et al*, 2015). However, such approaches largely depend on the types of p53 mutations a tumor carries, which potentially limits the broad use of each strategy in clinic. Thus, it becomes very critical to identify common regulators of different p53 GOF mutants for effective treatment of cancers.

Protein translation can be carried out by both cap-dependent and -independent pathways. When cap-dependent translation is globally inhibited under conditions such as cellular stresses, cells can continuously synthesize full-length p53 protein and ΔNp53 isoform by utilizing two IRES elements residing in the 5’UTR (hereafter termed IRES1) and the coding region (hereafter termed IRES2) of p53 mRNA, respectively (Candeias *et al*, 2006, Ray *et al*, 2006, Yang *et al*, 2006). The importance of IRES-dependent p53 protein production is further manifested in unstressed cells (Weingarten-Gabbay *et al*, 2014). A few proteins, including polypyrimidine tract binding protein (PTB), translation initiation factor DAP5, HDMX and HDM2, have been identified as putative IRES *trans*-acting factors (ITAFs) that preferentially bind to IRES2 of p53 mRNA (Grover *et al*, 2008, Malbert-Colas *et al*, 2014, Weingarten-Gabbay *et al*, 2014). To date, however, proteins that preferentially bind to p53 IRES1 remain unidentified. In the present study, we identified EZH2 as a p53 mRNA binding protein. We demonstrated that EZH2 bound to IRES1 of p53 mRNA and enhanced p53 protein translation in a methyltransferase-independent manner. We further showed that inhibition of such function of EZH2 induced synthetic lethality in p53 GOF-driven cancer cells.

## RESULTS

### RIP-seq analysis identifies EZH2 as a p53 mRNA-binding protein

Increasing evidence suggests that interactions with long non-coding RNAs (lncRNAs), such as *HOTAIR* and *XIST*, are important for the PcD activity of EZH2 (Rinn *et al*, 2007, Zhao *et al*, 2008). To define previously unrecognized oncogenic functions of EZH2, we sought to identify new EZH2-interacting RNAs in cancer cells. A previous study determined that EZH2 nonselectively binds to RNAs, at least under in vitro conditions while findings from other studies suggest that the PRC2 complex as a whole may not do the same in live cells (*Cifuentes-Rojas et al, 2014*, Davidovich *et al*, 2013). Given that the crosslink-based RIP is highly susceptible to contamination with non-specific RNAs (Kaneko *et al*, 2014), we performed native EZH2 RIP-seq in two prostate cancer cell lines without using a crosslink strategy. In addition to binding with lncRNAs, EZH2 selectively bound to a subset of messenger RNAs (mRNAs) encoding proteins highly relevant to cancer such as p53 (Figures 1A, 1B and Table EV1). Using RIP-coupled quantitative polymerase chain reaction (RIP-qPCR) we confirmed the selective association of the EZH2 protein with mRNAs of p53 and a few other genes such as *AKT1* and *KDM1A*, but not genes like *SKP2* (Figures 1C and EV1A-EV1D). These data indicate that EZH2 protein selectively binds to mRNAs of a subset of cancer-relevant genes including *TP53* in cells.

**Figure 1.**
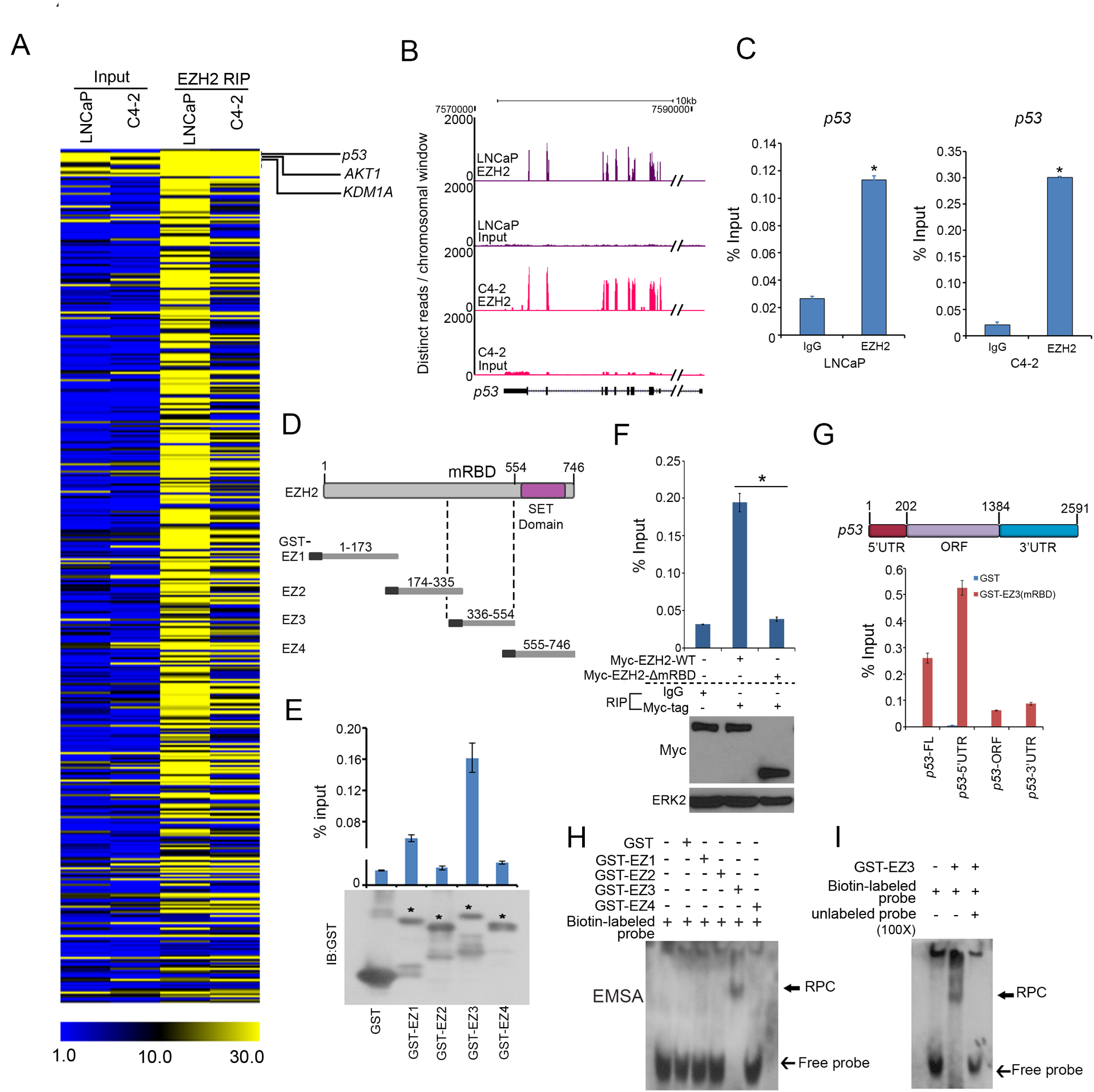
EZH2 binds to 5’UTR of *p53* mRNA. (A) Heat map showing mRNAs of genes immunoprecipitated by anti-EZH2 antibody in LNCaP and C4-2 prostate cancer cell lines, which was generated according to the distinct reads on chromosomal window from RIP-seq. (B) Screen shots from the UCSC genome browser showing signal profiles of *p53* mRNA immunoprecipitated by anti-EZH2 antibody in LNCaP and C4-2 cells. (C) RT-qPCR measurement of *p53* mRNA immunoprecipitated by IgG or anti-EZH2 antibody in LNCaP and C4-2 cells. Data shown as means ± SD (n=3). * *P*<0.01 comparing to IgG. (D) Schematic diagram of four GST-EZH2 recombinant proteins (EZ1 to EZ4). (E) Top, RT-qPCR analysis of *p53* mRNA in C4-2 cell lysate pulled down by GST or GST-EZH2 recombinant proteins EZ1-EZ4. Bottom, western blotting analysis of GST or GST-EZH2 proteins used for GST pull-down assay. Asterisks indicate the protein bands at expected molecular weight. (F) C4-2 cells were transfected with Myc-tagged EZH2-WT or EZH2-∆mRBD for 24 h and cells were harvested for RIP with IgG or anti-Myc tag antibody. Transfected proteins and pull-down *p53* mRNAs were analyzed by western blot and RT-qPCR, respectively. Data shown are means ± SD (n=3). ERK2, a loading control. (G) GST pull-down assay using in vitro transcribed different fragments of *p53* mRNA and indicated GST proteins followed by RT-qPCR analysis of pull-down *p53* mRNA. FL, full length; ORF, open reading frame; UTR, untranslated region. (H and I) EMSA evaluation of EZh2 binding of *p53* mRNA. GST-EZH2 recombinant proteins (EZ1 to EZ4) were incubated with biotin-labeled in vitro transcribed p53 5’UTR (biotin-labeled probe) in the presence or absence of 100-fold unlabeled p53 5’UTR (unlabeled probe), followed by PAGE and immune blotting with HPR-conjugated streptavidin.

### EZH2 binds to IRES1 of p53 mRNA

The p53 protein is a cellular gatekeeper that plays essential roles in maintaining genomic integrity and regulating cell growth, survival, and energy metabolism (Levine, 1997, Li *et al*, 2012). We chose to further characterize the molecular basis of the interaction between EZH2 protein and p53 mRNA and the biological consequences. We first examined which region of EZH2 protein is responsible for p53 mRNA binding. Glutathione S-transferase (GST)-EZH2 recombinant proteins were purified from bacteria as described (Chen *et al*, 2010) (Figure 1D) and incubated with lysate of LNCaP cells which express endogenous wild-type p53. GST pull-down products were subjected to RNA purification and reverse transcription quantitative PCR (RT-qPCR) analysis. The central region (amino acids 336-554), but not other parts of EZH2 protein bound strongly to p53 mRNA, and this region is therefore named as the p53 mRNA-binding domain (mRBD) (Figures 1D and 1E). RIP assays demonstrated that deletion of mRBD abolished EZH2 binding of p53 mRNA (Figure 1F). In vitro RNA binding assay showed that the mRBD in EZH2 bound primarily to the 5’UTR, but not the coding region and the 3’UTR of p53 mRNA (Figure 1G). These data suggest that EZH2 binds directly to p53 mRNA 5’UTR. To further validate this observation, we performed in vitro RNA electrophoretic mobility shift assay (EMSA) using purified human EZH2 and biotin-labeled p53 5’UTR as a probe. Consistent with GST pull-down results (Figures 1E and 1F), GST-EZ3 (mRBD), but not GST alone or other GST-EZH2 recombinant proteins formed an RNA-protein complex (RPC) with the p53 5’UTR (Figure 1H). The binding was dose-dependent and blocked by excessive amount of unlabeled p53 5’UTR (Figures 1I and EV2A), confirming that the interaction between EZH2 and p53 mRNA 5’UTR is specific. Together, these data suggest that EZH2 protein directly binds to the 5’UTR of p53 mRNA.

### EZH2 enhances IRES1-mediated p53 protein translation

Consistent with the in vitro RNA binding assay results (Figure 1G), the reciprocal biotin-labeled RNA pull-down assays demonstrated that endogenous EZH2 protein from LNCaP cell lysate were bound strongly by p53 mRNA 5’UTR, but very weakly by the 3’UTR and ORF (Figure 2A). As a positive control, EZH2 was readily pulled down by the HOTAIR lncRNA (Figure 2A). We further showed that a 120-nucleotide (nt) region (-120 to -1 nt) immediately adjacent to the translation start in the 5’UTR of p53 mRNA is critical for EZH2 binding (Figure 2B). Notably, this region almost completely overlaps with IRES1 (-130 to -1 nt) reported previously (Ray *et al*, 2006, Yang *et al*, 2006).

**Figure 2.**
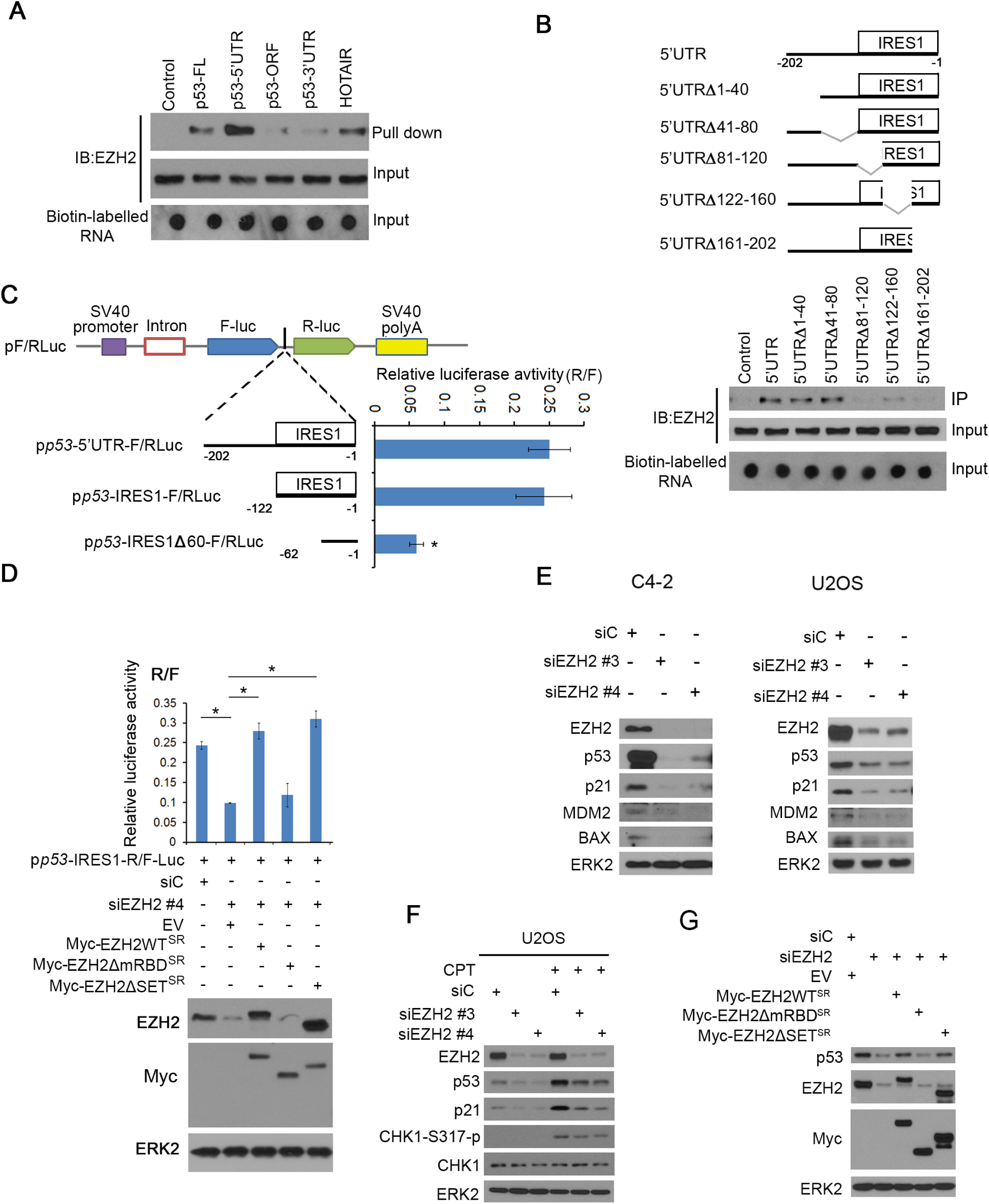
EZH2 upregulates p53 expression via binding to the IRES domain in 5’UTR. (A) Biotin pull down assay using biotin-labelled different fragments of p53 mRNA and HOTAIR (positive control) and C4-2 cell lysate followed by western blot with EZH2 antibody. (B) Biotin pull down assay as in (A) using different 5’UTR of p53 mRNA with different internal deletion mutants. Top, diagram of different p53 5’UTR deletion mutants. (C) Left, the linear map of the pRF bicistronic report plasmid. The SV40 promoter (purple box) was used to drive Firefly luciferase (Fluc) and Renilla luciferase (Rluc) gene transcription. Different p53 5’UTR fragments were inserted between the Fluc and Rluc genes. Right, at 24 h after transfection, cells were lysed and luciferase activities were measured using a dual-luciferase kit and the ratio of Fluc/Rluc was calculated. Data shown as means±SD (n=3). * *P*<0.01. (D) EZH2 knockdown C4-2 cells were transfected with the bicistronic reporter vector in combination with empty vector, Myc-tagged EZH2 WT or deletion mutants followed by western blot analysis with indicated antibodies (bottom) and luciferase assay (top). Data shown as means±SD (n=3). * *P*<0.01. (E) C4-2 and U2OS cell lines were transfected with non-specific control (siC) or two independent EZH2-specific siRNAs and harvested for western blot analysis with indicated antibodies. ERK2, a loading control. (F) U2OS cells were transfected with control (siC) or EZH2-specific siRNAs and treated with 200 nM of CPT followed by western blot analysis for indicated proteins. (G) C4-2 cells were transfected with control (siC) or EZH2-specific siRNAs and plasmids for empty vector, EZH2 WT or deletion mutants followed by western blot analysis for indicated proteins.

We further employed a dual reporter assay to determine whether EZH2 regulates p53 protein translation by binding to IRES1. We generated a series of luciferase reporter constructs by cloning different portions of the p53 5’UTR into a bicistronic plasmid (Figures 2C and EV2B). Similar to the previous reports in MCF7 and HeLa cells (Ray *et al*, 2006, Yang *et al*, 2006), both the entire 5’UTR and the IRES1 region of p53 mRNA exhibited translation-promoting activity in comparison to the empty vector (Figure 2C). However, deletion of 60 nt in the 5’ end of IRES1 (IRES1Δ60) largely diminished the activity of IRES1 (Figures 2C). Knockdown of endogenous EZH2 by small interference RNAs (siRNAs) significantly reduced the luciferase activity of the IRES-F/RLuc construct, and this effect was completely reversed by restored expression of siRNA-resistant wild-type EZH2 (EZH2-WT^SR^), SET-domain deletion (methyltransferase-deficient) mutant (EZH2ΔSET^SR^), but not the mRBD-deletion mutant (EZH2ΔmRBD^SR^) (Figure 2D). These data indicate that EZH2 enhances IRES1-dependent translation of p53 mRNA and this effect is mediated through the RNA-binding function, but not the methyltransferase activity of EZH2.

We also determined the effect of EZH2 on the steady-state level of p53 protein under physiological conditions. We knocked down EZH2 using two independent siRNAs in the C4-2 prostate cancer cell line and U2OS osteosarcoma cell line, both of which express wild-type p53. Knockdown of endogenous EZH2 decreased expression of endogenous p53 proteins in both cell lines and p53 downstream targets p21^CIP1^, MDM2 and BAX at both protein and mRNA levels (Figures 2E, EV2C and EV2D). Thus, EZH2 regulates expression of p53 protein and its downstream genes in unstressed cells.

p53 remains at low activity under normal conditions and becomes highly activated in response to genotoxic stresses. We treated U2OS cells with camptothecin (CPT), a chemotherapeutic drug that inhibits the re-ligation activity of topoisomerase-1 and therefore causes DNA double-strand breaks. CPT treatment increased expression of p53 protein and the downstream target p21^CIP1^ in control siRNA-treated cells (Figure 2F). The effectiveness of CPT was evident by increased phosphorylation of CHK1, a DNA damage checkpoint kinase (Figure 2F). Most importantly, CPT-induced upregulation of p53 protein expression was largely diminished by EZH2 knockdown (Figure 2F). These results suggest that EZH2 also regulates p53 protein expression in genotoxically stressed cells.

To determine whether EZH2 augments p53 protein expression via mRBD, we performed knockdown and rescue experiments. EZH2 knockdown-induced downregulation of p53 protein was almost completely reversed by restored expression of siRNA-resistant EZH2 (EZH2-WT^SR^) in C4-2 cells (Figure 2G). Similar results were obtained by restored expression of EZH2ΔSET^SR^, but not EZH2ΔmRBD^SR^ mutant (Figure 2G). These data suggest that EZH2 enhances p53 protein expression in cells in a manner dependent on binding to p53 mRNA.

The effect of EZH2ΔmRBD demonstrated in Figure 2G also indicates that EZH2 regulated p53 protein levels in a manner independent of the methyltransferase activity. To further confirm this observation, we treated C4-2 cells with the EZH2 enzymatic inhibitor GSK126 (McCabe *et al*, 2012b). As expected, GSK126 treatment effectively increased expression of EZH2-repressed PcD genes (e.g. *DAB2IP* and *BRACHYURY* (Wang *et al*, 2013a)) and decreased expression of EZH2-activated PcI genes (e.g. *TEME48*, *CKS2* and *KIAA0101* (Xu *et al*, 2012)), but it had no overt effect on p53 mRNA and protein expression (Figures EV3A-EV3C). Taken together, these data suggest that the function to bind to p53 mRNA, but not the methyltransferase activity is important for EZH2 to increase p53 protein expression in cells.

### EZH2 interacts with eIF4G and eIF4A and regulates p53 mRNA binding with polysomes

Without the requirement of eIF4E, both viral and cellular IRESs can mediate cap-independent protein translation through non-canonical binding with the eIF4G-eIF4A complex (Jackson *et al*, 2010, Komar & Hatzoglou, 2011, Pestova *et al*, 2001). We performed unbiased tandem affinity purification and mass spectrometry (TAP-MS) in 293T cells transfected with an empty vector containing S, Flag, and Biotin-binding-protein-(streptavidin)-binding-peptide (SFB) tags or SFB-tagged EZH2. The core components of PRC2, including SUZ12, EED and JARID2, were effectively pulled down by EZH2 (Figure 3A and Table EV2), indicating that the purification was successful. Importantly, eIF factors involved in cap-independent translation including eIF4G, eIF4A and eIF3 as well as poly(A)-binding protein-1 (PABP1) were also immunoprecipitated by SFB-EZH2 (Figure 3A). In contrast, the cap-binding protein eIF4E was not detected in the EZH2-associated protein complexes. The interaction of endogenous or Myc-tagged EZH2 with eIF4G2 and PABP1 was confirmed by reciprocal co-immunoprecipitation assays (Figures 3B and EV3D). Based upon these findings and those shown in Figures 1 and 2, we hypothesized that EZH2 promotes unconventional translation of p53 by binding to IRES1 in p53 mRNA 5’UTR and components of the cap-independent translation machinery proteins such as eIF4G and eIF4A (Figure 3C). To test this hypothesis, we performed polysome profiling in C4-2 cells with or without EZH2 knockdown. EZH2 knockdown resulted in approximately 50% reduction of p53 mRNA presented in polysomes (Figures 3D and EV3E). To determine whether this effect was specifically mediated by EZH2, we performed rescue experiments similar to the one shown in Figure 2G. Restored expression of EZH2 WT^SR^ and EZH2ΔSET^SR^, but not EZH2ΔmRBD^SR^ mutant was able to restore the level of p53 mRNA in polysomes (Figure 3E). We provide evidence that EZH2 regulates p53 mRNA association with polysomes in a methyltransferase-independent fashion.

**Figure 3.**
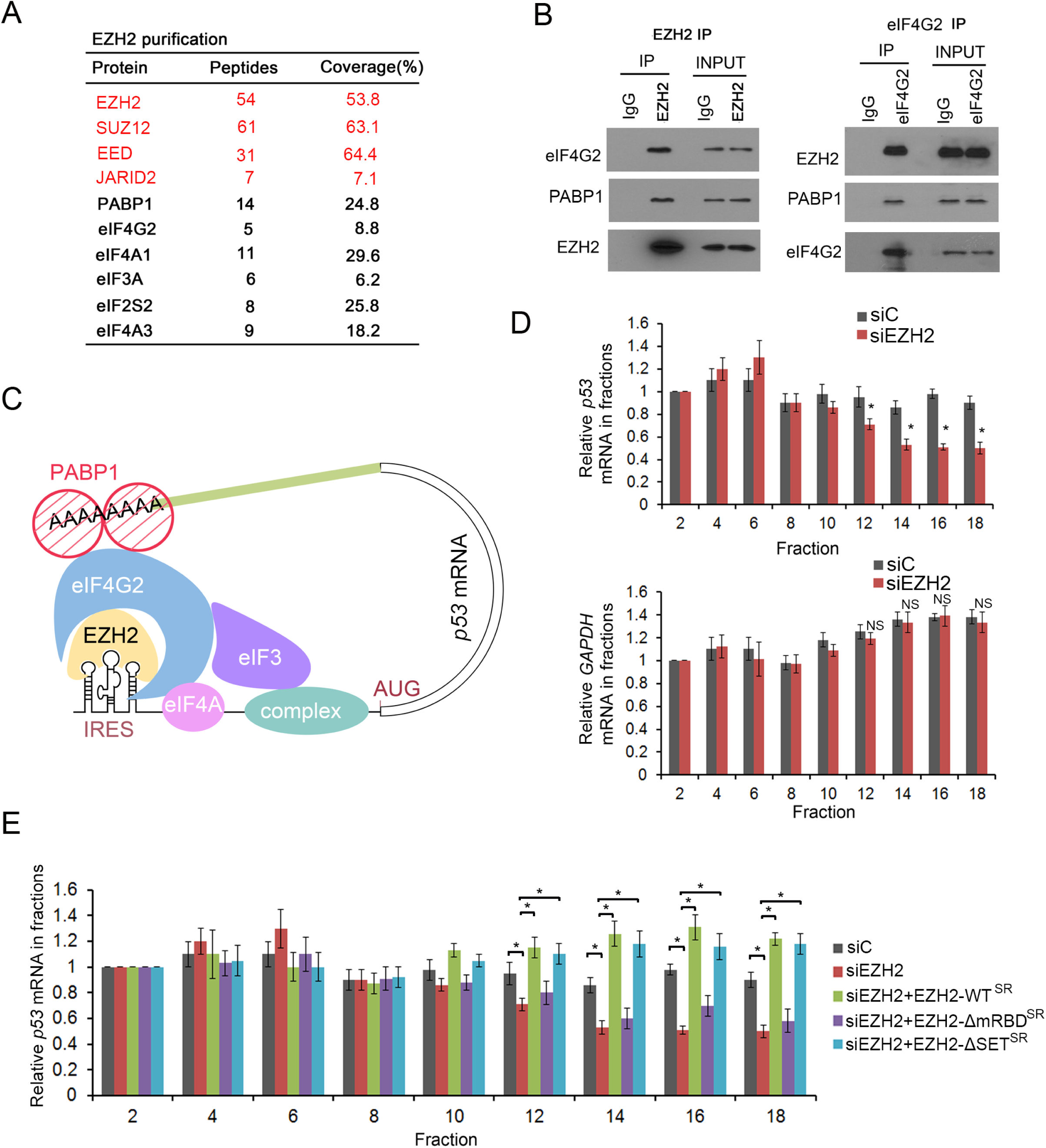
EZH2 binds with cap-independent translation complex and increases p53 protein translation. (A) List of top hits of proteins bound by EZH2 identified by TAP-MS. The number of peptides and coverage of each protein are included. (B) Reciprocal co-IP of endogenous EZH2 and eIF4G2 or PABP1 in C4-2 cells. (C) Schematic diagram depicting a hypothetical model wherein EZH2 augments p53 mRNA translation by binding with the IRES element in the 5’UTR of p53 mRNA and the cap-independent translation complex. (D) C4-2 cells were transfected control or (siC) or EZH2-specific siRNA and lysed for polysome fractionation. RNA was extracted from even-number fractions followed by RT-qPCR analysis for p53 and GAPDH mRNA. The β-ACTIN mRNA was used as an internal control. Data shown as means±SD (n=3). * *P*<0.01; NS, no significance. (E) C4-2 cells were transfected with control (siC) or EZH2-specific siRNAs in combination with empty vector, EZH2 WT or different deletion mutants followed by polysome fractionation and RT-qPCR. The β-ACTIN mRNA was used as an internal control. Data shown as means±SD (n=3). * *P*<0.01.

### EZH2 increases p53 mRNA stability

Increasing evidence suggests that the eIF4G-PABP1 interaction promotes the formation of a ‘closed’ mRNA loop that not only enhances ribosomal recruitment but also prevents mRNA decay. We sought to determine whether EZH2 regulates p53 mRNA stability. We examined the effect of EZH2 on the steady-state level of p53 mRNA. Knockdown of endogenous EZH2 by two independent siRNAs invariably decreased mRNA levels of endogenous wild-type p53 in both C4-2 and U2OS cell lines (Figure EV3F). This effect was completely reversed by restored expression of siRNA-resistant EZH2-WT^SR^ and EZH2ΔSET^SR^, but not the EZH2ΔmRBD^SR^ mutant (Figures 2D and EV3G). By measuring the rate of p53 mRNA decay, we demonstrated EZH2 knockdown shortened the half-life of p53 mRNA in both C4-2 and U2OS cell lines (Figure EV3H). Consistent with the finding that EZH2 did not bind to SKP2 mRNA (Figure EV1C), EZH2 knockdown had no overt effect on the half-life of SKP2 mRNA in these two cell lines (Figure EV3I). These results suggest that in addition to regulating p53 protein translation, EZH2 also enhances p53 expression by increasing mRNA stability.

### Ezh2 knockout decreases p53 mRNA and protein levels in a prostate cancer mouse model

Nucleotide sequence comparison showed that there was approximately 70% sequence similarity between human p53 mRNA 5’UTR and mouse and rat counterparts (Figure EV4A). Similar to the human homolog, the predicted secondary structure of rodent p53 mRNA 5’UTRs also exhibits three major potential stem-loop moieties (Figure EV4B). In agreement with these observations, RIP-qPCR assay showed that Ezh2 also associated with p53 mRNA in murine cell lines (Figure EV4C). *PTEN* is a tumor suppressor gene that is frequently mutated or deleted in human prostate cancers, and homozygous deletion of the *Pten* gene invariably promotes tumorigenesis in the mouse prostate. Homozygous deletion of *Pten* induces upregulation of p53 in prostate cancers in mice (Chen *et al*, 2005), but the underlying mechanism remains poorly understood. Several studies independently show that *Pten* knockout upregulates Ezh2 mRNA and proteins in mouse prostate tumors (Ding *et al*, 2014, Kuzmichev *et al*, 2005, Mulholland *et al*, 2011). We therefore employed the *Pten* knockout mouse model to determine whether EZH2 plays a causal role in the regulation of p53 expression in vivo. We examined whether increased expression of Ezh2 correlates with p53 expression in prostate tumors in this model. As expected, homozygous *Pten* deletion resulted in upregulation of both p53 and Ezh2 proteins in prostate tumors in mice (Figures 4A and 4B). Importantly, Ezh2 levels highly correlated with p53 protein expression in a cohort of *Pten*-knockout mouse prostate tumors (Figure 4C). To determine the causal role of Ezh2 in p53 upregulation in *Pten*-null mouse prostate tumors, we generated prostate-specific *Pten*;*Ezh2* double knockout mice. *Ezh2* deletion decreased expression of p53 protein and mRNA as well as its downstream gene *Bax* in the background of *Pten* deletion (Figures 4A, 4D and 4E). Thus, similar to the effect in human cancer cell lines cultured in vitro, Ezh2 also plays a causal role in the regulation of p53 mRNA and protein expression under in vivo conditions.

**Figure 4.**
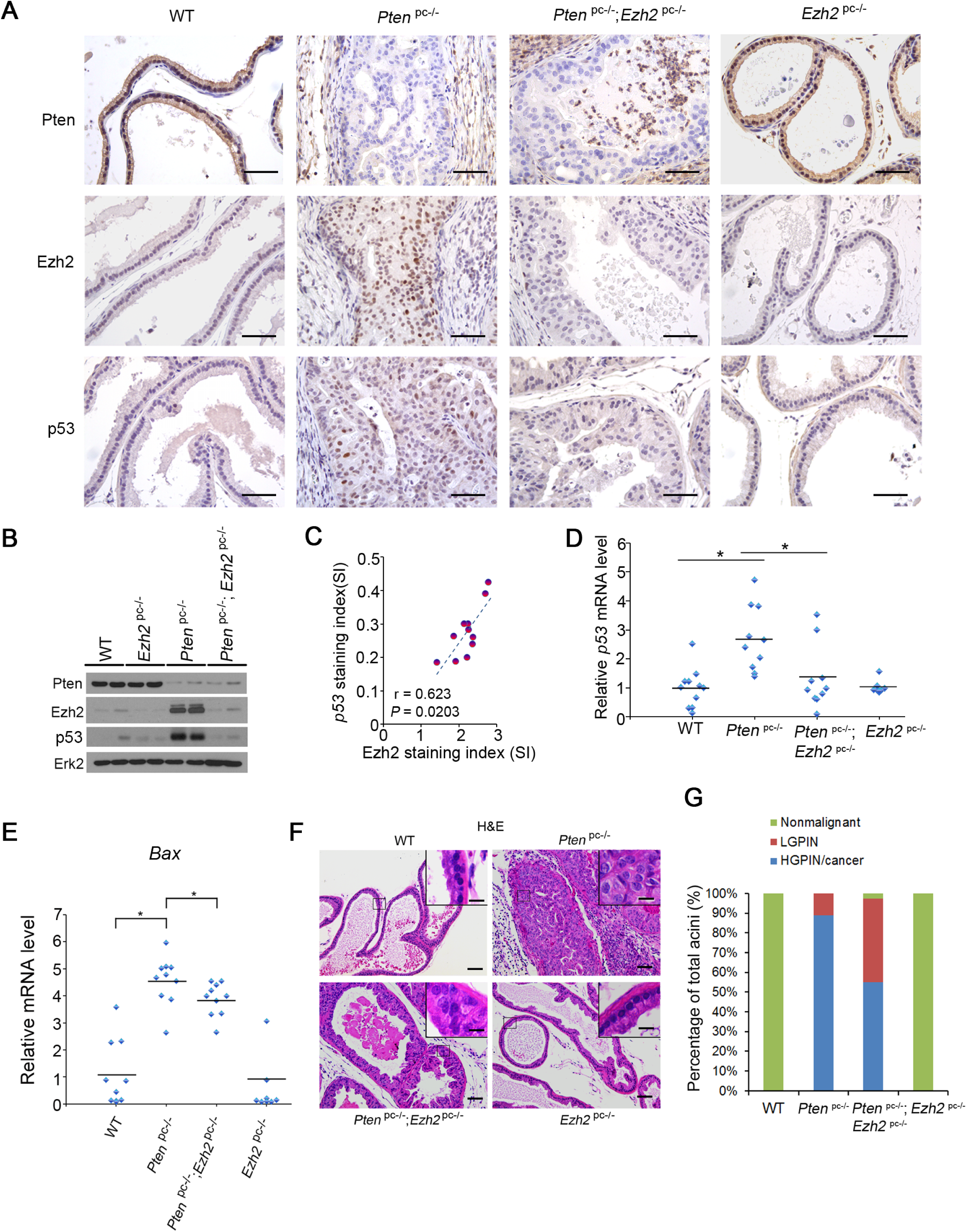
*Ezh2* knockout decreases p53 expression in the *Pten*-null prostate cancer mouse model. (A) IHC analysis of Pten, Ezh2 and p53 proteins in prostatic tissues of 4-month-old mice with different genotypes: *Pten*^pc-/-^ (n=11); *Ezh2*^pc-/-^ (n=8); *Pten*^pc-/-^;*Ezh2*^pc-/-^ (n=10) and ‘wild-type’ littermate controls (n=10). Scale bar, 50 µm. (B) Western blot analysis of expression of indicated proteins in four different groups of mice as described in (A) (n=2 mice/group). (C) Analysis of correlation of Ezh2 and p53 protein levels determined by IHC in *Pten* knockout (*Pten*^pc-/-^) mice (n=11). See IHC scoring details in Methods section. (D) RT-qPCR analysis of p53 mRNA expression in four different genotypes of mice as described in (A). * *P*<0.01. (E) RT-qPCR analysis of mRNA expression of p53 downstream target gene *Bax* in four different genotypes of mice as indicated. * *P*< 0.01. (F) H&E analysis of ventral prostate (VP) of 4-month-old mice with indicated genotypes. The inset shows a high magnification image of the representative (framed) area in each panel. Scale bar, 50 µm. Scale bar in inset, 10 µm. (G) Quantification of nonmalignant, low-grade PIN (LGPIN) and high-grade PIN (HGPIN) or cancerous acini in the prostates (including anterior prostate (AP), ventral prostate (VP) and dorsolateral prostate (DLP)) of mice with the indicated genotypes and number as in (A).

Consistent with the previous report that EZH2 level is extremely low in normal human prostatic tissues (Varambally *et al*, 2002), we demonstrated that Ezh2 protein was barely detectable in the prostate of wild-type mice (Figures 4A and 4B). It is not surprising that homozygous deletion of *Ezh2* in prostatic epithelium had little or no effect on p53 mRNA and protein expression and prostate epithelium morphogenesis (Figures 4A, 4B, 4D, 4F and 4G). Following *Pten* deletion, even though Ezh2 protein levels were markedly elevated in *Pten*-knockout prostate tumors (Figures 4A and 4B), there were a significant portion of acini remained at the high grade prostatic intraepithelial neoplasia (HGPIN)/cancer stage after co-deletion of *Ezh2* in the *Pten*-deleted tumors (Figures 4F and 4G). These results are consistent with the finding that the levels of p53 mRNA and protein and its downstream target gene *Bax* was significantly downregulated in *Pten;Ezh2* double knockout tumors compared with *Pten* single knockout tumors (Figures 4D and 4E). Thus, depletion of EZH2 in malignant tissues that express high levels of EZH2 results in undesirable downregulation of wild-type p53, which may play a favorable role in promoting tumorigenesis. These findings also provide a plausible explanation for the phenomenon that homozygous deletion of *Ezh2* failed to completely block *Pten* deletion-induced tumorigenesis in the prostate with the background of wild-type p53.

### Expression of EZH2 and p53 positively correlates in human cancers

The finding that EZH2 increases p53 protein levels by enhancing mRNA stability and protein translation in cultured human cancer cells and mouse tumors prompted us to determine the correlation of these two proteins in cancer patient specimens. Meta-analyses of previously published gene expression profiling data revealed a positive correlation between EZH2 and p53 mRNA in a variety of human cancer types examined, including prostate, brain, colorectal cancer, and sarcoma (Figures EV5A-EV5H). *EZH2* and *p53* mRNA levels also correlated in prostate cancers of The Cancer Genome Atlas (TCGA) cohort (Figure 5A). The correlation between EZH2 and p53 was further confirmed at both mRNA and protein levels by RT-qPCR and IHC, respectively in prostate cancer specimens from two independent cohorts of patients (Figures 5B-5D). IHC analysis also showed that EZH2 protein expression correlated with p53 protein levels in various other cancer types, including brain, colorectal, pancreatic cancer, and sarcoma (Figures 5E and 5F). These results indicate that EZH2 and p53 expression are correlated positively at both mRNA and protein levels in many types of human cancer.

**Figure 5.**
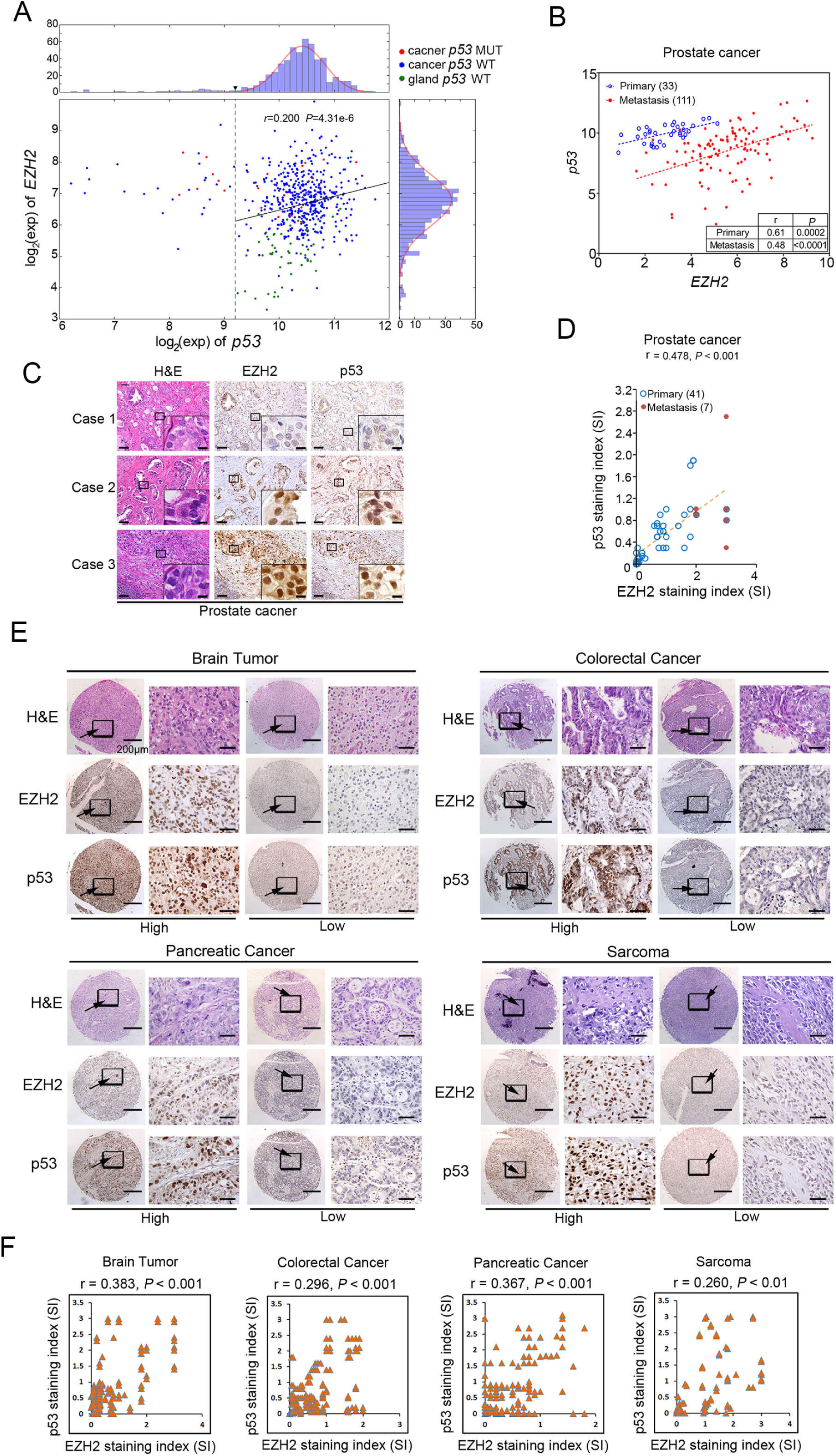
EZH2 correlates with p53 expression in various types of human cancer. (A) Correlation analysis of EZH2 and p53 mRNA levels in primary prostate cancer specimens of the TCGA cohort. (B) Correlation analysis of EZH2 and p53 mRNA expression measured by RT-qPCR in primary and metastatic prostate cancer specimens of a University of Washington cohort. (C) IHC analysis of EZH2 and p53 protein expression in human prostate cancer tissues of a Mayo Clinic cohort. Case 1 and 2 show low and high expression in primary tissues, respectively; case 3 shows high expression in lymph node metastasis. Scale bar, 50 µm. Scale bar in inset, 10 µm. (D) Correlation analysis of EZH2 and p53 protein expression in primary and metastatic prostate cancer specimens of a Mayo Clinic cohort. (E) IHC analysis of EZH2 and p53 protein expression in commercially purchased TMA with patient specimens of brain, colorectal, pancreatic cancer and sarcoma. Scale bar in low magnification image, 200 µm; Scale bar in high magnification image, 50 µm. (F) IHC data as shown in (E) were quantified and used for correlation analysis of EZH2 and p53 protein expression in patient specimens of brain, colorectal, pancreatic cancer and sarcoma. Pearson correlation r and *P* values in each cancer type are indicated.

### EZH2 increases the level of mutated p53 mRNA and protein in cancer cells

It is generally accepted that mutated p53 protein is much easier to be detected by IHC in cancer specimens than the unmutated counterpart, and therefore detection of high-level p53 protein expression is often utilized as a proxy for the presence of p53 mutations (Muller & Vousden, 2014). A previous association study reveals that EZH2 overexpression positively correlates with the high level expression of p53 protein in esophagus squamous cell carcinomas, many of which were shown to express mutated p53, although the molecular mechanism underlying the correlation was not explored (Yamada *et al*, 2011). Since both wild-type and mutated p53 mRNAs share the same IRES1 in the 5’UTR, we sought to determine whether EZH2 modulates the expression of mutated p53 in a manner similar to the wild-type counterpart. Knockdown of EZH2 invariably decreased the steady-state level of different forms of p53 mutants at both protein and mRNA levels in various cancer cell lines of different tissue origins, including VCaP (R248W) and DU145 (P223L and V274F p53 mutants) prostate cancer cell lines and U251 (R273H p53 mutant) and T98 (M237I p53 mutant) glioblastoma cell lines (Figures EV5I-EV5L). These data indicate that EZH2 also regulates mutated p53 expression in cancer cells.

### EZH2 cooperates with p53 GOF mutants to promote cancer growth and metastasis

Most p53 mutations in human cancer are missense mutations and often cluster at a few “hotspot” amino acids (Muller & Vousden, 2014). Previous studies in cell culture, xenograft and genetically engineered mouse models suggest that some of the “hotspot” mutations, such as R175H, R248W and R273H, are GOF mutations that gain new functions in promoting cancer invasion, metastasis, and progression (Dittmer *et al*, 1993, Freed-Pastor *et al*, 2012, Olive *et al*, 2004, Weissmueller *et al*, 2014). We sought to determine whether the cancer-driven functions of p53 GOF mutants are regulated by EZH2. VCaP cells express R248W GOF mutant of p53.

Knockdown of R248W mutant by shRNAs largely decreased VCaP cell growth in 2D and 3D culture conditions and in mice (Figures 6A-6D, EV5M and EV5N). Importantly, ectopic expression of EZH2ΔSET, but not the p53 mRNA-binding-deficient mutant EZH2ΔmRBD significantly increased growth of VCaP cells in culture and in mice, and no such effect was detected in R248W p53 mutant knockdown cells (Figures 6A-6D, EV5M and EV5N). Thus, EZH2 augments p53 GOF mutant-mediated cancer growth in a methyltransferase-independent manner.

**Figure 6.**
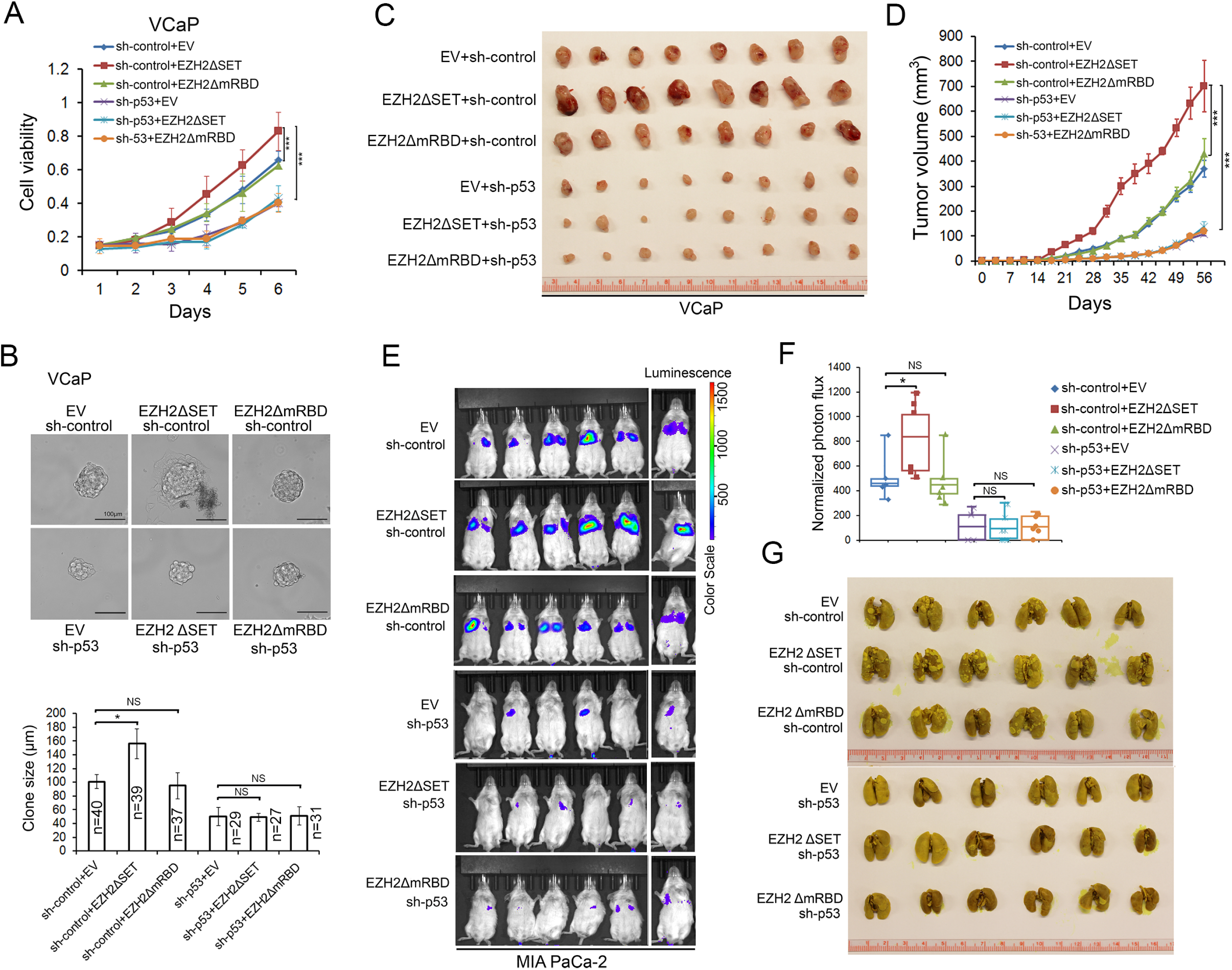
EZH2 enhances p53 GOF mutant-mediated cancer growth and metastasis independently of its methyltransferase activity. (A and B) VCaP cells were transfected with control (sh-control) or p53-specific shRNA and shRNA-expressing stable cells were infected with lentivirus for empty vector (EV) or deletion mutants of EZH2. Cell growth in 2D (A) and 3D (B) conditions were determined by MTS assay and measurement of clone size, respectively. *** *P* < 0.001; NS, no significance. (C and D) VCaP cells (1×10^7^) infected with lentivirus as in (A) were injected subcutaneously into NSG mice (n = 8/group). Tumors were measured by caliper twice a week. Data are shown as means ± SD (D), and tumors at the end point of measurement were isolated and photographed (C). Statistical significance was determined by two-tailed Student’s *t*-test for tumors at day 56. *** *P* < 0.001. (E-G) Luciferase-expressing MIA PaCa-2 cells (2×10^6^) infected with lentivirus as in (A) were injected via tail vein into NSG mice (n = 6/group). At 12 weeks after injection, mice were subjected to bioluminescent imaging, and images were recorded (E) and bioluminescent signals were quantified (F). Lungs were isolated from mice, stained with Bouin’s solution, and photographed (G). The white spots on lungs (stained in yellow) are metastatic tumors. * *P* < 0.05; NS, no significance.

p53 GOF mutants can also drive cancer metastasis (Yue *et al*, 2015). We sought to determine to what extent EZH2 regulation of p53 GOF mutant expression contributes to cancer cell invasion and metastasis. Ectopic expression of full-length mRNA of p53 R273H and R248W mutants largely enhanced invasion of p53-null PC-3 prostate cancer cells (Figures EV6A-EV6F). EZH2 knockdown not only decreased protein levels of these mutants, but also substantially inhibited cell invasion augmented by p53 R248W and R273H (Figures EV6A-EV6F). MIA PaCa-2 is a highly metastatic pancreatic cancer cell line which expresses an endogenous p53 GOF mutant R248W. Expression of EZH2ΔSET mutant not only elevated R248W protein levels, but also largely increased cell invasion, and no such effects were observed for the p53 mRNA-binding-deficient mutant EZH2ΔmRBD (Figures EV6G and EV6H). EZH2ΔSET-induced cell invasion was completely abolished by knockdown of endogenous p53 R248W mutant (Figures EV6G and EV6H). These data indicate that EZH2 enhances p53 GOF mutant-induced cell invasion.

We further examined whether EZH2 promotes cancer metastasis through interaction with p53 GOF mutant mRNA. To this end, we generated MIA PaCa-2 cells expressing the luciferase reporter gene. Consistent with the results of cell invasion assay (Figures EV6G and EV6H), ectopic expression of EZH2ΔSET mutant, but not the p53 mRNA-binding-deficient mutant EZH2ΔmRBD largely increased MIA PaCa-2 cell metastasis to lung (Figures 6E-6G and EV6I). Most importantly, knockdown of endogenous p53 R248W mutant almost completely abolished EZH2ΔSET mutant-induced lung metastasis of MIA PaCa-2 cells in mice (Figures 6E-6G and EV6I).

*SHARP1* and *CCNG2* are two cancer metastasis-associated genes reported as the targets of mutated p53 (Adorno *et al*, 2009). Expression of these two genes was upregulated by the methyltransferase-deficient mutant EZH2ΔSET, but not the p53 mRNA-binding-deficient mutant EZH2ΔmRBD in control knockdown MIA PaCa-2 cells (Figure EV6J). However, this effect was completely abolished by knockdown of endogenous p53 mutant R248W (Figure EV6J). These data suggest that EZH2 binding of p53 mRNA is important for p53 GOF mutant-mediated cancer cell invasion and metastasis, implying that EZH2 is a viable target for treatment of tumors harboring p53 GOF mutations.

### EZH2 depletion induces synthetic vulnerability in p53 GOF mutant-expressing cancer cells

Based on the data described above, we hypothesized that in cancer cells expressing WT p53, EZH2-enhanced expression of p53 acts against EZH2-mediated oncogenesis (Figure 7A, left). In contrast, in cancer cells expressing p53 GOF mutants, EZH2 and mutated p53 work cooperatively in favor of cancer progression (Figure 7A, right). In support of this hypothesis, analysis of TCGA prostate cancer data showed that high levels of EZH2 proteins significantly correlate with the worse biochemical recurrence-free survival only in patients expressing mutated p53, but not p53 WT or loss (Figure 7B). These data suggest that EZH2 represents a promising therapeutic target for p53-mutated cancer cells, especially those with GOF mutants. To test this notion, we measured cell viability and growth of cancer cells expressing either R248W GOF mutant or p53 WT in the presence or absence of EZh2 inhibition. We treated VCaP (p53 GOF mutant R248W) and C4-2 (p53 WT) cells with different EZH2 inhibitory agents, including GSK126 (inhibits EZH2 methyltransferase enzymatic activity) (McCabe *et al*, 2012b), DZNep (inhibits EZH2 methyltransferase enzymatic activity and EZH2 protein expression) (Tan *et al*, 2007), and EZH2 antisense oligonucleotides (ASOs) (inhibits EZH2 protein expression). As expected, treatment of these agents invariably decreased H3K27me3 levels, increased expression of PcD gene *DAP2IP* and downregulated expression of PcI genes *CEP76*, *RAD51C* and *TEME48* in both cell lines (Figures 7C and EV6K). However, treatment with DZNep and ASO, but not GSK126 also decreased expression of WT p53 and its downstream target p21^CIP1^, a cell growth inhibitory protein in C4-2 cells (Figure 7C). No such effect on p21^CIP1^ expression was observed in R248W-expressing VCaP cells (Figure 7C). Accordingly, treatment with DZNep and ASO resulted in much greater inhibitory effect on viability of VCaP cells cultured in 2D and 3D compared to C4-2 cells (Figures 7D and EV7A), and similar results were obtained from colony formation assays (Figure EV7B). Together, these data show that depletion of EZH2 protein expression, but not the inhibition of its enzymatic activity induces synthetic vulnerability in p53 GOF mutant-expressing cancer cells.

**Figure 7.**
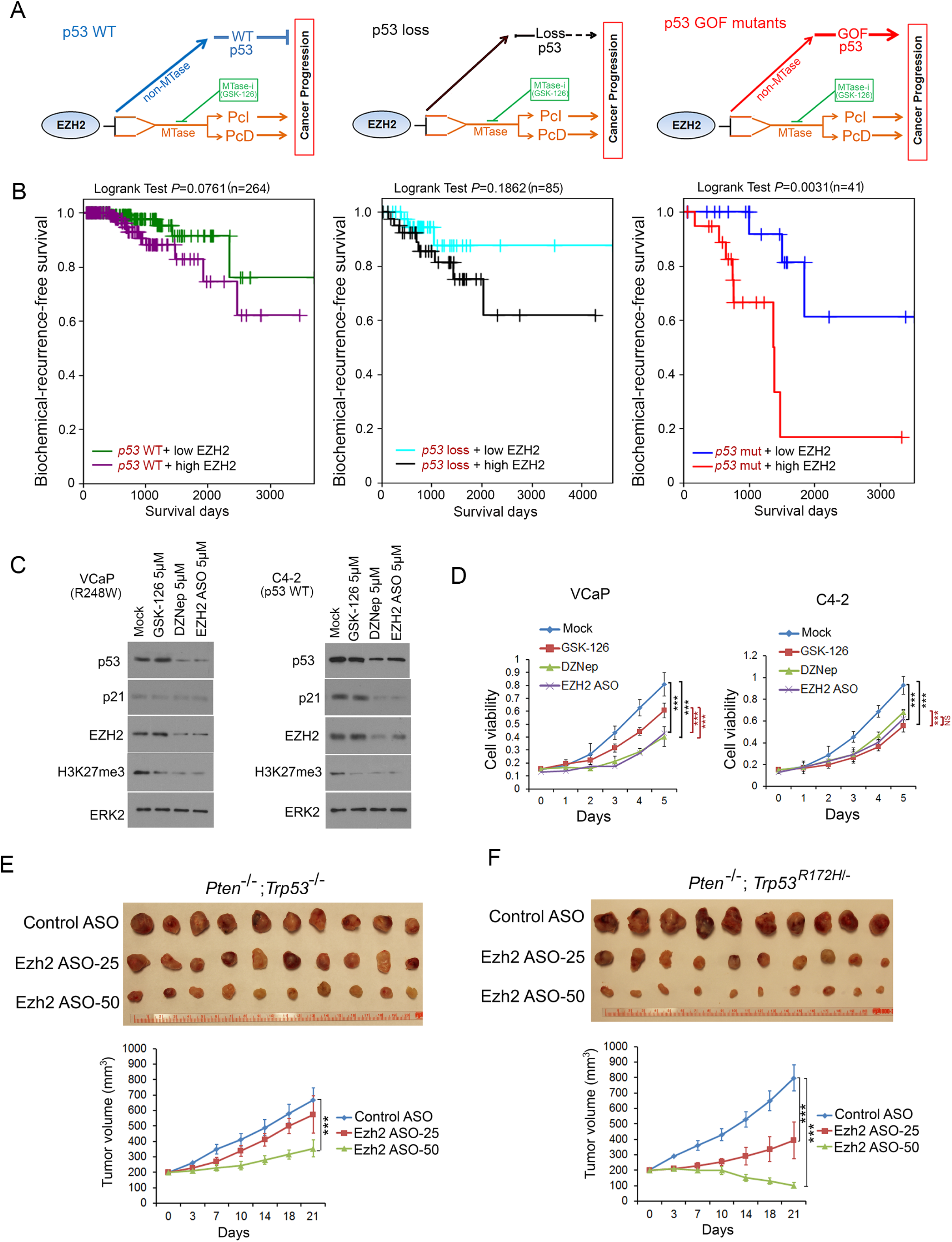
Depletion of EZH2 expression inhibits growth of p53 GOF-mutated cancer cells. (A) Schematic diagram depicting the distinctive impacts of the functional interplay between EZH2 protein and p53 mRNA on progression of cancers expressing WT p53, no p53 or GOF mutated p53. MTase, methyltransferase. (B) Kaplan-Meier plots showing the association of EZH2 overexpression with biochemical recurrence of prostate cancer in patients from the TCGA cohort. The EZH2 was separated into two groups (high or low expression); p53 was separated into 3 groups (wild type, loss and mutated). (C and D) VCaP and C4-2 cells were treated with GSK-126, DZNep, and EZH2 ASO, followed by western blots with indicated antibodies (C) and MTT assay (D). *** *P* < 0.001. (E and F) Effect of Ezh2 ASOs on growth of murine prostate cancer allografts. Mice (n=10) bearing the *Pten*^−/-^;*Trp53*^−/-^ (E) and *Pten*^−/-^;*Trp53*^R172H/-^ allografts (F) were treated according the scheme shown in (Figure S7D) with control ASOs (50 mg kg^−1^) or Ezh2 ASOs at different doses (25 mg kg^−1^ and 50 mg kg^−1^). Tumor growth was measured twice a week for 3 weeks and the data are shown in the bottom panels. Each allograft was isolated by the end of ASO treatment and photographed (top panels). Statistical significance was determined by two-tailed Student’s *t*-test. *** *P* < 0.001.

### EZH2 depletion inhibits growth of p53 GOF mutant-expressing tumors in vivo

To determine the inhibitory effect of EZH2 depletion on growth of p53 GOF mutant-expressing tumors in mice, we generated murine prostate cancer cell lines from prostate-specific *Pten*^−/-^;*Trp53*^−/-^ and *Pten*^−/-^;*Trp53*^R172H/-^ mutant mice (Figure EV7C). These tumor cells were inoculated subcutaneously into NSG mice. When the size of allografts reached up to 200 mm^3^, mice were randomly grouped and treated intraperitoneally (i.p.) with control or Ezh2-specific ASOs twice a week for 3 weeks (Figure EV7D). We demonstrated that volumes of *Pten*^−/-^;*Trp53*^−/-^ tumors were slightly decreased in mice treated with 25 mg of Ezh2 ASO (Ezh2 ASO-25) but significantly decreased after treated with 50 mg of Ezh2 ASO (Ezh2 ASO-50) in comparison to control ASO treatment (Figures 7E and 7F). Most importantly, p53 GOF mutant-expressing tumors (*Pten*^−/-^;*Trp53*^R172H/-^) were much more sensitive to Ezh2 ASO treatment compared to p53-deficient (*Pten*^−/-^;*Trp53*^−/-^) tumors (Figures 7E and 7F). Notably, little or no effect of Ezh2 ASO treatment on mouse weight loss was detected (Figure EV7E). RT-qPCR analysis indicated that *Ezh2* mRNA expression was effectively depleted by Ezh2 ASO treatment, especially at the high dose in the tumors and so was true for *Trp53* mRNA expression (Figures EV7F and EV7G). In contrast, only modest reduction in *Ezh2* mRNA levels was observed in some normal tissues such as brain and testis, but not in other tissues examined including heart, intestine, kidney, liver, lung and muscle and generally the alterations were not statistically significant with two exceptional occasions (Figure EV7F). Importantly, no significant changes were observed in *Trp53* mRNA levels in all the normal tissues examined and this was presumably due to the fact that Ezh2 levels were much lower in normal tissues than that in tumors (Figures EV7F and EV7G). This data is consistent with our findings in *Ezh2* knockout prostates (Figures 4A, 4B and 4D), implying that negligible level of EZH2 may have limited leverage in regulation of p53 expression in normal tissues. Together, these data suggest that depletion of EZH2 protein with strategies such as ASOs can be a viable therapeutic arsenal for treatment of advanced cancer, especially those expressing p53 GOF mutants.

## DISCUSSION

### Non-methyltransferase function of EZH2 in cancer

PcD and PcI are two major functions of EZH2 identified thus far. Since both roles are dependent on the methyltransferase activity and implicated in cancer development and progression (Varambally *et al*, 2002, Xu *et al*, 2012), a significant amount of effort has been put into developing EZH2 enzymatic inhibitors. A few such inhibitors including GSK126 and EPZ-6438 have been developed and are currently in phase I clinical trials for treatment of B-cell lymphomas and advanced solid tumors (Knutson *et al*, 2014, McCabe *et al*, 2012b, Vaswani *et al*, 2016). To our knowledge, however, no favorable reports have been documented yet. In the present study, we provide evidence that EZH2 binds to IRES1 in the 5’UTR of both wild-type and mutated *p53* mRNA and increases mRNA stability and cap-independent protein translation. We further show that this function of EZH2 is independent of its methyltransferase activity. Thus, we identify a previously unrecognized non-methyltransferase function of EZH2 in cancer. Our findings also suggest that targeting EZH2 expression rather than targeting its enzymatic activity could be a more effective therapeutic option, particularly in tumors expressing GOF mutated p53.

### Mechanistic explanation of the dichotomous role of EZH2 in cancer

Paradoxical roles of EZH2 and PRC2 have been seen in different types of human cancer. It is well established that EZH2 plays an oncogenic role and correlates with the progressiveness or stages in most types of solid cancer. However, deletions of *EZH2* and other PcG genes such as *SUZ12* occur in a subset of hematopoietic malignancies such as T-cell acute lymphoblastic leukemia (T-ALL) (Ntziachristos *et al*, 2012). Our discovery of the positive regulation of p53 mRNA and protein by EZH2 supports the model wherein EZH2 may exert dichotomous roles in oncogenesis of cells expressing a wild-type p53 (Figure 7A, left). We also provide evidence that in tumors harboring p53 GOF mutant, EZH2 upregulates and works cooperatively with mutated p53, thereby favoring cancer progression (Figure 7A, right). Moreover, similar to our findings in solid tumors (Figure 7B), the prognosis of leukemia patients with a mutation in the *TP53* gene is worse than those expressing a wild-type p53 (Peller & Rotter, 2003). This model is further supported by a previous report that EZH2 overexpression and p53 mutations frequently occur in late-stage cancers (Varambally *et al*, 2002). Thus, our findings not only provide a plausible explanation for the dichotomous roles of EZH2 occurring in certain cancer types, such as a subset of hematopoietic malignancies, but also reveal a biological and mechanistic basis for the co-occurrence of EZH2 overexpression and p53 mutations in many advanced cancer types.

### EZH2 binding of mRNAs of cancer relevant genes

EZH2 is a known RNA-binding protein. By binding to noncoding RNAs (ncRNAs) such as XIST, RepA and HOTAIR, EZH2 has been shown to work together with other PcG proteins to promote X chromosome inactivation, developmental patterning, and maintenance of stem cell pluripotency (Plath *et al*, 2003, Rinn *et al*, 2007, Wang *et al*, 2013b, Zhao *et al*, 2008). Binding of EZH2 with ncRNAs such as HOTAIR and MALAT1 also promotes cancer growth and metastasis (Wang *et al*, 2015, Wang *et al*, 2011). Therefore, the biological and clinical significance of EZH2 interaction with ncRNAs have been extensively studied. Using the unbiased RNA-seq approach, in the present study we demonstrated that EZH2 binds to a group of message RNAs which code functionally important proteins such as p53. Most importantly, we provide evidence that EZH2 binding of p53 mRNA is functional. Specifically, we showed that EZH2 increases p53 mRNA stability and promotes cap-independent protein translation of p53 mRNA by binding to the IRES element in the 5’UTR. Thus, our finding of EZH2 binding of message RNAs largely broadens our understanding of the functional significance of RNA binding of EZH2, thereby representing a framework for the discovery of new roles of EZH2 in developmental and cancer biology.

### A new strategy to inhibit p53 GOF mutants in cancer

Given that mutated p53 proteins are generally considered less druggable (Lehmann & Pietenpol, 2012), endeavor has been largely focused on targeting the downstream effectors of each individual GOF mutant of p53 (Adorno *et al*, 2009, Weissmueller *et al*, 2014, Zhu *et al*, 2015). We demonstrated that different from the inhibition of EZH2 enzymatic activity, depletion of EZH2 protein expression with ASOs selectively suppresses growth of cancer cells expressing GOF mutated p53, highlighting that targeting EZH2 induces synthetic vulnerability in p53 GOF mutated cancer cells. While we found that EZH2 regulates mRNA and protein levels in cultured cancer cells, we provided evidence that homozygous deletion of *Ezh2* in normal tissues (e.g. normal prostate gland) or treatment with *Ezh2* ASOs had no drastic impact on *Trp53* mRNA and protein expression in mice. In agreement with these observations, we also show that EZH2 expression is much lower in normal tissues than that in tumors and therefore, it is not surprising that negligible EZH2 expression level may have limited leverage in regulation of p53 expression in normal tissues. These findings stress that targeting the upstream regulator such as EZH2 offers a new opportunity to inhibit various p53 GOF mutants with diversified functions in cancer, and such treatment appears to have little or no harm on normal tissues, which is fully supported by our observation that Ezh2 ASO treatment almost had no effect on body weight of mice. Thus, our findings not only indicates EZH2 is a viable therapeutic target in p53-mutated cancer, but also suggest that inhibition of functionally diversified p53 GOF mutants is achievable by single targeting of EZH2 as a common upstream regulator, which therefore represents a new paradigm of targeted therapy of p53-mutated cancer.

## STAR METHODS

Detailed methods are provided in the supplementary information.

## ACKNOWLEDGEMENTS

We thank the patients and their families for their altruism in participating in research studies. We also thank Xinbin Chen, Zhenbang Chen, and Da-Qing Yang for reagents and suggestions, Wenqian Hu from Mayo Clinic for providing facilities for polysome fractionation, Youngsoo Kim and Robert MacLeod from Ionis Pharmaceuticals Inc for providing EZH2 ASOs, members of Huang laboratory for their constructive comments for the study, Colm Morrissey, Robert Vessella, Larry True, Xiaotun Zhang and all other members of the University of Washington rapid autopsy team for their tremendous efforts. This work was supported in part by grants from the National Institutes of Health (CA134514, CA130908 and CA193239 to H.H.), the Department of Defense (W81XWH-09-1-622 and W81XWH-14-1-0486 to H.H. and W81XWH-17-1-0415 TO P.S.N.), the Mayo Clinic Center for Individualized Medicine (to H.H.), the Pacific Northwest SPORE in Prostate Cancer P50CA097186 (to P.S.N.) and the Prostate Cancer Foundation (to P.S.N.).

